# ALPK1 and TIFA dependent innate immune response triggered by the Helicobacter pylori type IV secretion system

**DOI:** 10.1101/139998

**Authors:** Stephanie Zimmermann, Lennart Pfannkuch, Munir A. Al-Zeer, Sina Bartfeld, Manuel Koch, Jianping Liu, Cindy Rechner, Meike Soerensen, Olga Sokolova, Alla Zamyatina, Paul Kosma, André P. Mäurer, Frithjof Glowinski, Klaus-Peter Pleissner, Monika Schmid, Volker Brinkmann, Michael Naumann, Marion Rother, Nikolaus Machuy, Thomas F. Meyer

**Affiliations:** Department of Molecular Biology, Max-Planck Institute for Infection Biology, 10117 Berlin, Germany; Institute of Experimental Internal Medicine, Otto von Guericke University Magdeburg, 39120 Magdeburg, Germany; Department of Chemistry, University of Natural Resources and Life Sciences-Vienna, 1190 Vienna, Austria; Steinbeis Innovation, Center for Systems Biomedicine, 14612 Berlin-Falkensee, Germany; Research Center for Infectious Diseases, ZINF, Institute for Molecular Infection Biology, IMIB, University of Würzburg, 97080 Würzburg, Germany

**Keywords:** *Helicobacter pylori*, NF-κB signaling, genome wide RNAi screen, innate immune response, inflammation, alpha kinase 1 (ALPK1), TRAF interacting protein with forkhead associated domain (TIFA), D-*glycero*-β-D-*manno*-heptose 1,7-bisphosphate (HBP), Type IV secretion system (T4SS)

## Abstract

Activation of transcription factor NF-κB is a hallmark of infection with the gastric pathogen *Helicobacter pylori* and associated with inflammation and carcinogenesis. Genome-wide RNAi screening revealed numerous hits involved in *H. pylori*-, but not IL-1β- and TNF-α- dependent NF-κB regulation. Pathway analysis including CRISPR/Cas9-knockout and recombinant protein technology, immunofluorescence microscopy, immunoblotting, mass spectrometry and mutant *H. pylori* strains, identified the *H. pylori* metabolite D-*glycero-β*-D-*manno-heptose* 1,7-bisphosphate (βHBP) as a cagPAI type IV secretion system (T4SS)-dependent effector of NF-κB activation in infected cells. Upon pathogen-host cell contact, TIFA forms large complexes (TIFAsomes) including interacting host factors, such as TRAF2. NF-κB activation, TIFA phosphorylation as well as TIFAsome formation depended on a functional ALPK1 kinase, highlighting the ALPK1-TIFA axis as core of a novel innate immune pathway. ALPK1-TIFA-mediated NF-κB activation was independent of CagA protein translocation, indicating that CagA translocation and HBP delivery to host cells are distinct features of the pathogen’s T4SS.

## Introduction

*H. pylori* chronically colonizes the gastric mucosa of about half the world’s population (Parkin, 2006). Infections can lead to ulcers, lymphoma of the mucosa-associated lymphoid tissue (MALT) and gastric adenocarcinoma (reviewed in (Salama et al., 2013)). It appears in two major strains, defined by the absence or presence of a type IV secretion system (T4SS), encoded by the *cag* pathogenicity island (cagPAI) that functions in the translocation of *H. pylori’s* only known effector protein, CagA, into host cells, where it is phosphorylated (Backert et al., 2000). The cagPAI positive strains exhibit particular strong inflammatory potential and increased pathogenicity (Blaser et al., 1995; Crabtree et al., 1991).

Activation of the transcription factor family nuclear factor kappa B (NF-κB) plays a central role in inflammation and carcinogenesis (DiDonato et al., 2012; Pasparakis, 2009). It can be triggered by various stimuli, such as the inflammatory cytokines tumor necrosis factor α (TNF-α), interleukin 1β (IL-1β), and infections by various pathogens. The NF-κB family is comprised of five members, including p65 (RelA), which in the inactivated state is sequestered in the cytoplasm by its inhibitor, inhibitor of kappa B alpha (IκBα). Upon activation, the IκB kinase (IKK) complex phosphorylates IκBα, which is subsequently degraded to release p65 for translocation into the nucleus, where it guides the transcription of target genes linked to inflammation and anti-apoptosis (Hacker and Karin, 2006; Hoffmann and Baltimore, 2006). Its dual function in activation of inflammation and cell survival makes NF-κB a lynchpin in disease development. These features are of particular importance in the gastrointestinal system, where chronic inflammation and tissue damage act as tumor promoters (Quante and Wang, 2008), but our understanding of the underlying regulatory network is still incomplete for many inducers. Several RNAi loss-of function studies have identified new factors involved in NF-κB pathway activation including stimuli of the nucleotide-binding oligomerization domain-containing protein (NOD)1 and NOD2 receptors (Bielig et al., 2014; Lipinski et al., 2012; Warner et al., 2013; Yeretssian et al., 2011), Epstein Barr virus (Gewurz et al., 2012), *Neisseria gonorrhoeae* (Gaudet et al., 2015), and *Shigella flexneri* infections (Milivojevic et al., 2017). These studies have shown that NF-κB signaling, although sharing core elements, is highly complex and diverse.

Here we report on a high content cell-based RNAi screening approach, addressing *H. pylori-induced* NF-κB activation. Amongst several host factors, we identified α-kinase 1 (ALPK1) and TRAF-interacting protein with FHA domain (TIFA) as key mediators of *H. pylori* induced NF-κB activation. Upon *H. pylori* infection, TIFA undergoes the formation of large protein complexes (TIFAsomes), including TRAF2 and additional host factors involved in NF-κB signaling. We demonstrate that the bacterial metabolite heptose-1,7-bisphosphate (HBP) triggers this particular route of NF-κB activation in a T4SS-dependent manner. Our findings reveal important mechanistic insight into the ability of the innate immune system to discriminate between less and highly virulent *H. pylori* traits (Koch et al., 2016) and the process of NF-κB activation that depends on the pathogen’s T4SS (Backert and Naumann, 2010).

## Results

### New regulators of *H. pylori*-induced NF-κB activation

As a read-out system for the identification of factors involved in NF-κB pathway activation we used nuclear translocation of a p65-GFP fusion protein which is amenable for analysis by automated microscopy (Bartfeld et al., 2010) (Figure 1A). Besides *H. pylori*, we used TNF-α and IL-1β as stimuli for NF-κB activation. Robustness of the system was assessed by targeting known factors of the NF-κB pathway. Accordingly, knockdown of TNF-α receptor 1 (TNFR1) and of the adaptor protein myeloid differentiation primary response protein 88 (MYD88) blocked the signals by TNFα and IL-1β, respectively. Moreover, combined IKK-α and IKK-β knock-down abolished activation by TNFα, IL-1β as well as by *H. pylori* (Figure 1B). Details of three screens performed are summarized in Figure S1A-C. The first screen using a human kinase siRNA library yielded 15 primary hits with increased responsiveness to *H. pylori* infection, 4 of which were confirmed by at least 2 independent siRNAs. These included 2 genes not previously implicated in NF-κB signaling: ALPK1 and CDC2-related kinase, arginine/serine rich (CRKRS) (Bartfeld, 2009) (Table S1). To extend our search, a second screen was carried out with only *H. pylori* infection as inducer, using two siRNA libraries targeting the whole and the druggable genome. This analysis yielded 347 hits: 235 positive and 112 negative regulators. Out of these, 200 of the positive and 100 of the negative regulators were further validated using additional siRNAs and all three inducers, which yielded 43 positive (21.5%) and 33 negative *H. pylori*-induced regulators (33.0%). Comparison to NF-κB induction by IL-1β or TNF-α indicated that 21 of the positive and 24 of the negative regulators were strongly biased to *H. pylori* as an inducer (Figure 1C, Table S2).

**Fig. 1:**
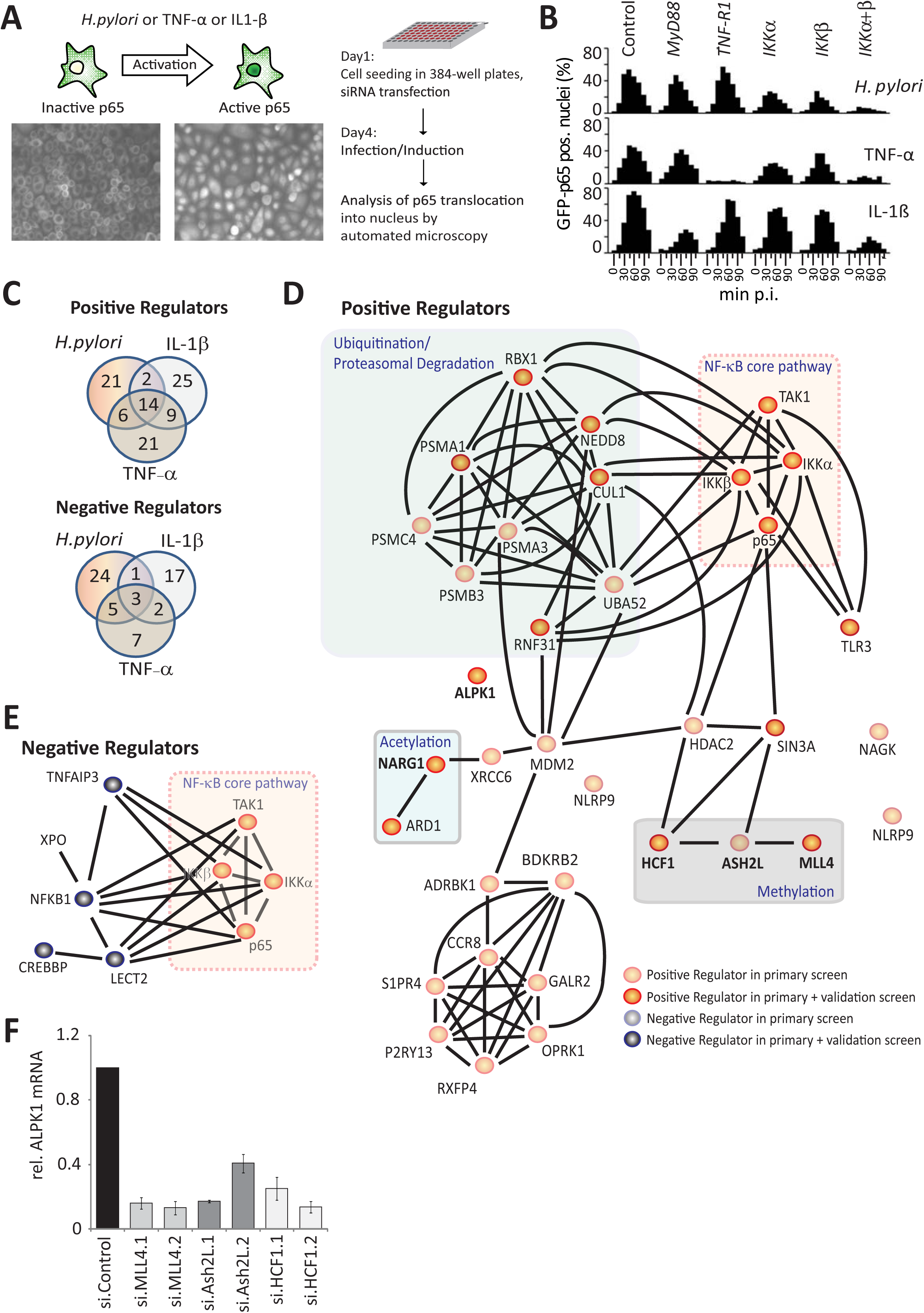
Genome-wide RNAi screen for *H.-pylori* induced NF-κB activation. (A) Screen setup: 72 h posttransfection with siRNAs AGS SIB02 cells (GFP-p65) were infected with *H. pylori* or induced with IL-1β or TNF-α, fixed and analyzed for p65 translocation into the nucleus. (B) Knockdown of NF-κB pathway activators leads to reduced p65 translocation in cells treated as in (A) and fixed at 0, 15, 30, 45, 60, 75, 90 or 105 min followed by quantification via automated microscopy. Inhibitory siRNAs used are IKKα and IKKβ for *H. pylori*, TNF-R1 for TNF-α or MyD88 for IL-1β. Control siRNA is Allstars. (C) Venn diagram showing the validated factors. Apart from factors functioning for all inducers, 21 positive and 24 negative regulators unique for the *H. pylori*-induced NF-κB pathway were found. (D, E) Selected protein-protein interactions among (D) positive or (E) negative regulators from the screen in connection to the NF-κB core pathway, identified with a confidence score of ≥0.4 from the STRING database. Hits found to be unique for *H. pylori* NF-κB signaling are shown in bold. Ubiquitination and proteasomal degradation is known to be heavily interconnected with the NF-κB core pathway. Additionally, we identified an acetylation complex formed by ARD1 and NARG1 and methylation complex consisting of HCF1, MLL4 and Ash2L to be important positive regulators in *H. pylori* NF-κB pathway. In contrast, the identified kinase ALPK1 is not connected to the NF-κB pathway. Among negative regulators we found the known NF-κB inhibitors TNFAIP3, CREBBP and NFKB1 (F) siRNA-mediated loss of MLL4, Ash2L or HCF1 strongly reduced the level of ALPK1 mRNA, as identified by qRT-PCR.

STRING analysis (Szklarczyk et al., 2011) of our datasets revealed prominent protein interaction networks (Figure 1D and 1E). Notably, one obvious sub-network was formed by known general NF-κB key regulators, consisting of IKK-α (Chuk), IKK-β (IKBKB), TAK1 (MAP3K7) and p65 (RelA) itself, corroborating the functional robustness of our approach (Figure 1D). The central roles of ubiquitination and proteasomal degradation in NF-κB signaling was well reflected by the data set (Oeckinghaus et al., 2011). Apart from major members of the proteasome (PSMA1, PSMB3, PSMA3 and PSMC4), we identified RBX1, NEDD8 and Cullin-1 as necessary for ubiquitination (Figure 1D). The RING finger-like domain-containing protein RBX1 is part of cullin-RING-based E3 ubiquitin-protein ligase (CRLs) complexes, which are regulated by NEDD8 ligation (Scott et al., 2011). We also identified RNF31 as a positive regulator for *H. pylori*, TNF-α and IL-1β NF-κB signaling. A prominent negative regulator, for example, included the deubiquitinase tumor necrosis factor alpha induced protein 3 (TNFAIP3), known to downregulate NF-κB (Harhaj and Dixit, 2011) (Figure 1E). Several of the identified hits, although not uniquely involved in *H. pylori*-induced signaling, had only a weak effect on TNF-α-induced and none in IL-1β-induced NF-κB signaling. Of these, ARD1 and NARG1, which form an acetyltransferase complex connected with the ubiquitinylation machinery (Arnesen et al., 2005), were further validated (Figure S1D). The regulators most specific for *H. pylori* also included α-kinase 1 (ALPK1) (Figure S1E, Table S2), an atypical kinase previously implicated in apical protein transport and phosphorylation of myosin IA (Heine et al., 2005). Confirmation by Western blotting showed that ALPK1 was required for degradation of IκBα after infection with *H. pylori* but not after stimulation with TNFα or IL-1β (Figure S1E). In addition, we included the TRAF-interacting protein with FHA domain (TIFA) in our validation rounds, as it had the highest score of any gene in our genome-wide RNAi screen, albeit for only one siRNA (Table S2).

### Upstream regulators of ALPK1

Notably, our analyses revealed a cluster of histone-modifiers as *H. pylori* specific hits (Figure 1D), including host cell factor C1 (HCF1), absent, small or homeotic 2 like (ASH2L), lysine K-specific methyltransferase 2B (MLL4), and SIN3 transcription regulator family member A (SIN3A) (Dou et al., 2005; Smith et al., 2005; Tyagi et al., 2007; Wysocka et al., 2003; Yokoyama et al., 2004). With HCF1 as a known co-regulator of transcription (Tyagi et al., 2007), we hypothesized that it may indirectly affect expression of other determinants detected in the screens. To our surprise, quantitative RT-PCR (qRT-PCR) guided by initial microarray analysis, indicated that ALPK1 expression was abrogated after HCF1 knockdown (Figure 1F). Similarly, knockdown of ASH2L or MLL4, known to act in concert with HCF1, turned out to abrogate ALPK1 expression as well (Figure 1F). Thus, methylation events mediated by the HCF1 methyltransferase complex appeared to be required for housekeeping expression of ALPK1 and the effects of the HCF1, ASH2L and MLL4 on *H. pylori*-induced NF-κB activation are likely caused indirectly via their influence on ALPK1 gene expression. These findings nominated ALPK1 as a central component of NF-κB signal transduction.

### Role of ALPK1 in NF-κB activation

To further consolidate the function of ALPK1 in *H. pylori*-specific NF-κB activation, an ALPK1 deficient cell line was generated using CRISPR/Cas9 technology (Figure S2A and S2B). Knockout of ALPK1 in several independent cell lines fully prevented p65 translocation after infection (Figure 2A), more strikingly than any of the siRNAs did. Yet, ALPK1 knockout did not affect the response to TNF-α or IL-1β (Figure 2B). Consistently, ALPK1 deficiency abrogated infection-induced TAK1 and IκB-α phosphorylation and subsequent IκB-α degradation but did not influence the response to TNF-α treatment (Figure 2C). Cells lacking ALPK1 were unable to mount an IL-8 response upon infection, as evidenced by significantly reduced IL-8 mRNA levels in infected ALPK1 deficient cells (Figure 2D). To test if p65 translocation could be rescued, ALPK1 deficient cells were transfected with recombinant Myc-Flag-ALPK1. Indeed, only rescued cells exhibited NF-κB activation upon *H. pylori* infection (Figure 2E, upper panel), whereas TNF-α treatment triggered p65 translocation also in ALPK1 deficient cells (Figure 2E, lower panel). These results identify ALPK1 as a strictly essential mediator of NF-κB activation in *H. pylori* infection.

**Fig. 2.**
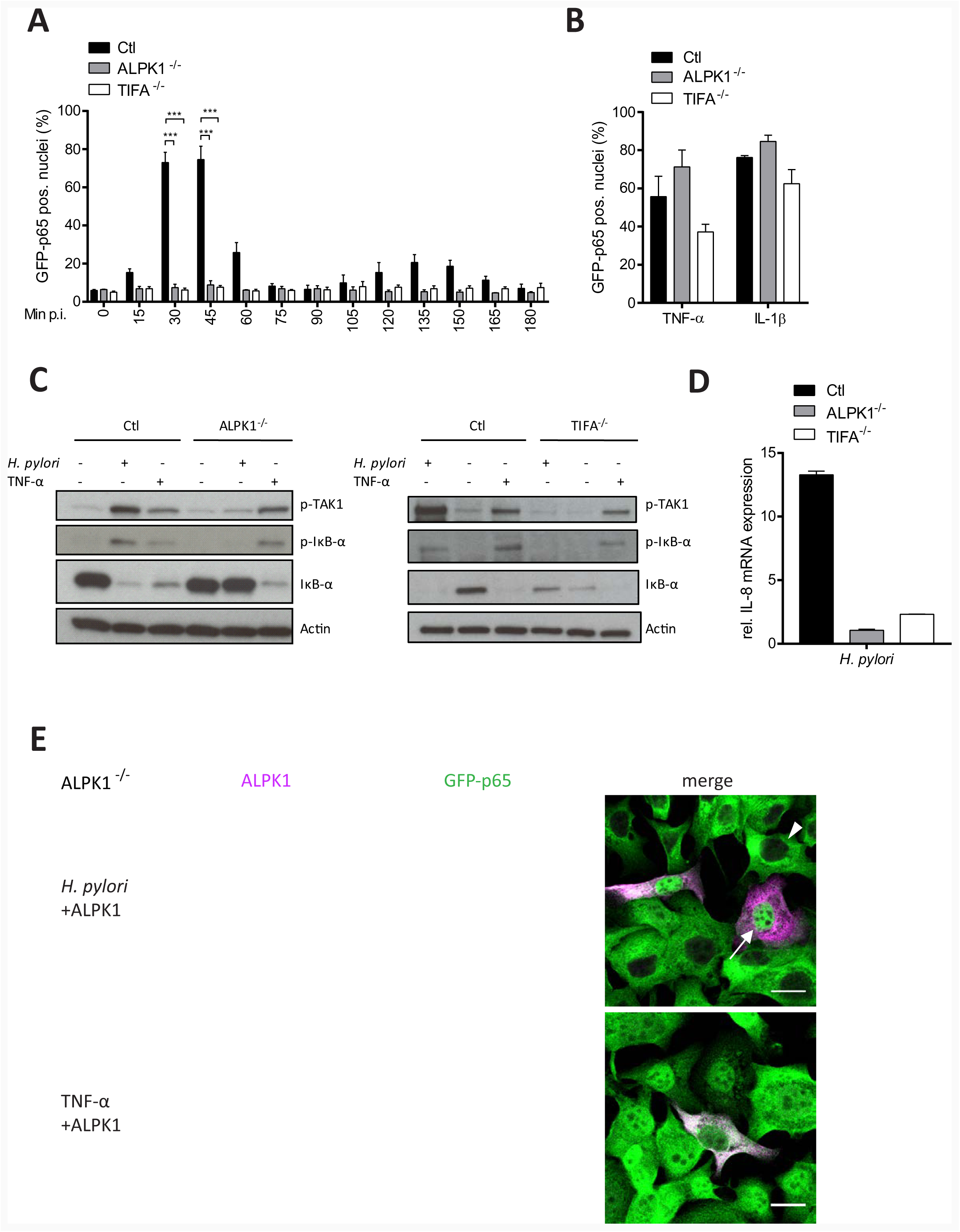
ALPK1 and TIFA are essential for the *H. pylori*-induced activation of NF-κB. CRISPR/Cas9-mediated silencing of ALPK1 or TIFA in AGS SIB02 cells (A) impedes NF-κB activation after infection with *H. pylori* MOI 100 for the indicated times. Shown is percentage of cells with nuclear p65 per well. (B) Silencing does not influence TNF-α- or IL-1β-induced NF-κB activation. (C) Loss of ALPK1 or TIFA blocks TAK1 phosphorylation at Thr-184/187 and IκB-α phosphorylation at Ser32. IκB-α degradation was blocked specifically in ALPK1 or TIFA deficient cells at 30 min post *H. pylori* infection at MOI 100. Silencing ALPK1 did not inhibit TNF-α-dependent (20 ng/ml for 30 min) NF-κB activation. Blot representative of at least two independent experiments. (D) qRT-PCR analysis of IL-8 transcription in ALPK1 and TIFA depleted AGS cells infected with *H. pylori* at MOI 100 for 30 min, normalized to noninfected control cells. (E) ALPK1 rescues NF-κB activation upon *H. pylori* infection (MOI 100 for 30 min) in ALPK1 depleted AGS SIB02 cells expressing GFP-p65 (green) transfected with Myc-Flag-ALPK1 for 24 h. Cells were labelled for Myc (magenta). ALPK1-expressing cells show nuclear translocation of p65 (top). Cells activated instead with TNF-α (20 ng/ml, for 30 min) also show nuclear translocation of p65. Arrowheads: non transfected, not activated cells, arrows: transfected, activated cells. Scale bar: 20 μm. Data in graphs represents mean ± SEM of three independent experiments. Statistical analysis was performed using Student’s t-test (*** *p* ≤ 0.0001).

### TIFA is crucial for *H. pylori-induced* NF-κB activation acting downstream of ALPK1

Simultaneously, we also tested CRISPR/Cas9-generated TIFA knockout cells to illuminate its role in *H. pylori*-specific NF-κB activation (Figure S2C and D). Interestingly, the TIFA knockout line failed to show any sign of *H. pylori*-specific NF-κB activation, even after extended monitoring for 3 h (Figure 2A), while p65 translocation readily occurred upon treatment with IL-1β or TNF-α (Figure 2B). Further, knockout of TIFA specifically abrogated *H. pylori*-induced phosphorylation of TAK1 and IκBα, as well as the subsequent degradation of IκBα (Figure 2C) but had no effect on TNFα-induced TAK1 and IκBα phosphorylation. Consequently, TIFA knockout cells did not show IL-8 mRNA expression upon infection, as measured by qRT-PCR 3 h p.i. (Figure 2D). Next, we sought to determine the cellular localization of TIFA by confocal immunofluorescence microscopy. Non-infected AGS SIB02 cells transfected with Myc-Flag-TIFA revealed diffuse TIFA fluorescence (red) throughout the cytoplasm and the nucleus whereas p65 location (green) was restricted to the cytoplasm and excluded from the nucleus (Figure S3A upper panel). Infection with wild type *H. pylori* or TNF-α treatment induced p65 nuclear translocation by 30 min (middle and lower panel). Consistent with NF-κB activation, only infection with wild type *H. pylori*, but not TNF-α treatment, induced the formation of so-called TIFAsomes (Gaudet and Gray-Owen, 2016) (Figure S3A). To further define the importance and specificity of TIFA oligomer formation upon infection, TIFA deficient cells were transfected with Myc-Flag-TIFA or a Myc-Flag-TIFA mutant for T9, an important tyrosine phosphorylation site for TIFA activation (Huang et al., 2012). In cells that remained non-transfected, only TNF-α treatment but not *H. pylori* infection induced p65 translocation (Figure 3A, arrow head upper panel and lower panel). Successful transfection with wild type, but not with the TIFA mutant T9A, restored both p65 translocation and TIFAsome formation upon *H. pylori* infection (arrow upper panel and middle panel). To monitor TIFAsome formation in more detail, we transfected TIFA deficient AGS cells with Tomato-TIFA and infected them with *H. pylori*. Time-lapse video microscopy of live cells confirmed infection-induced TIFAsome formation (from 7 min p.i.) and p65 translocation (from 12 min p.i.) only in transfected cells (Movie S1). TIFAsome formation could not be observed in ALPK1 deficient cells but was rescued by co-transfection with ALPK1 (Figure 3B) indicating that ALPK1 acts upstream of TIFA and thus plays a crucial role in the phosphorylation of TIFA T9 required for TIFAsome formation.

**Fig. 3:**
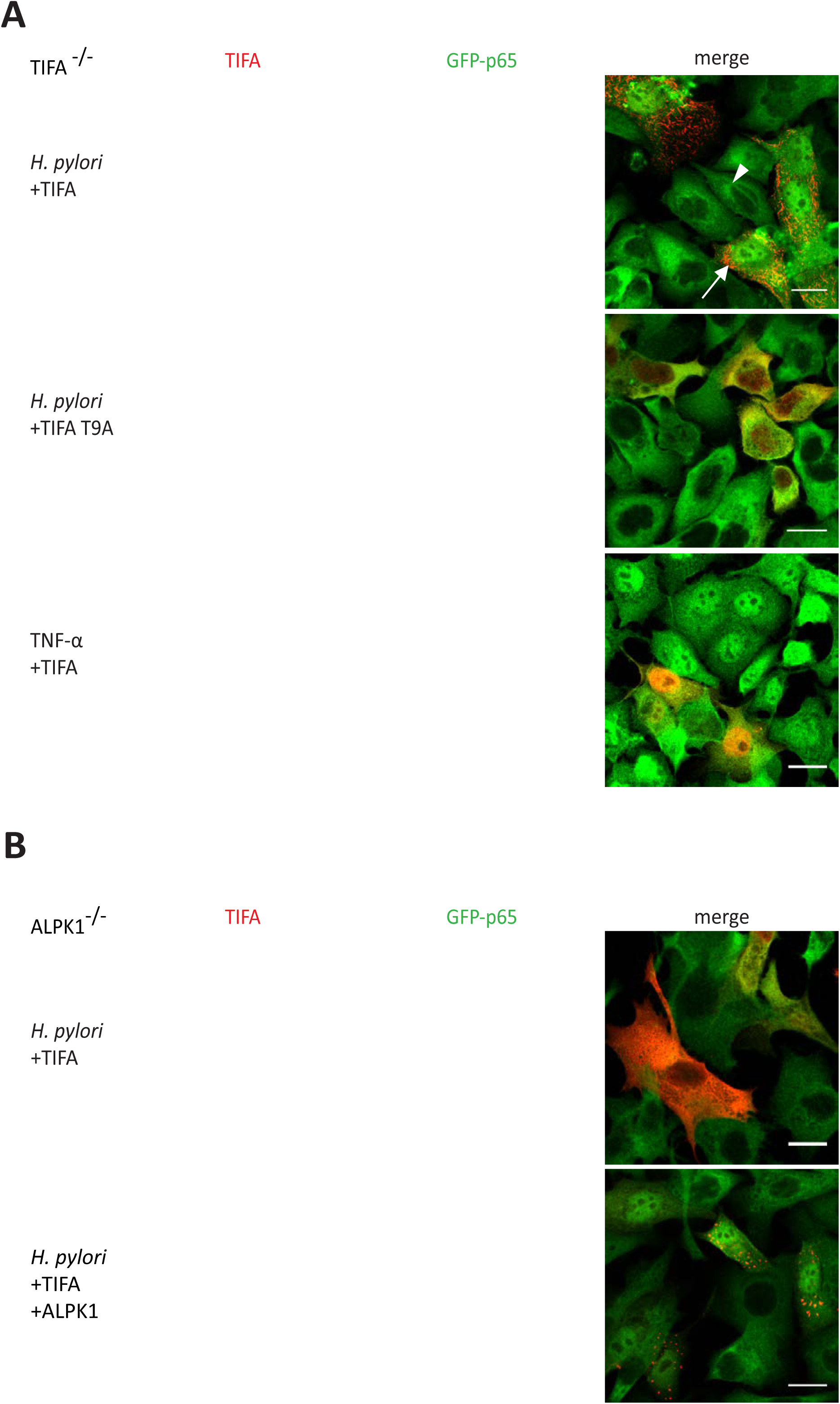
TIFA activates NF-κB upon *H. pylori* infection through formation of TIFAsomes dependent on ALPK1. (A) AGS SIB02 were depleted for TIFA using CRISPR/Cas9 and were transfected for 24 h with wild type or T9A mutated Myc-Flag-TIFA, or an empty vector cDNA constructs prior to infection. Only wild type TIFA rescues TIFAsome formation and NF-κB activation upon *H. pylori* infection (MOI 100, 30 min). Cells activated with TNF-α (20 ng/ml, 30 min) did not show TIFAsome formation. TIFA labelled in red. Arrowheads: non transfected, not activated cells, arrows: transfected, activated cells. Scale bar: 20 μm. (B) Lack of ALPK1 hinders TIFAsome formation in cells infected with *H. pylori* and can be rescued by transfection with ALPK1. AGS SIB02 were depleted for ALPK1 using CRISPR/Cas9 and transfected for 24 h with Myc-Flag-TIFA (red) or Tomato-TIFA (red) combined with Myc-Flag-ALPK1. *H. pylori* infection (MOI 100 for 30 min) did not induce TIFAsomes or NF-κB activation unless cells were co-transfected with ALPK1 and TIFA. Scale bar: 20 μm.

### TIFAsomes are multifactorial protein complexes

To identify protein–protein interactions (PPI) of TIFAsomes by mass spectrometry (MS) we used a Myc-Flag-TIFA fusion, which was transiently overexpressed in AGS cells for pull-down of TIFA associated proteins (Table S3). MS discovered a total of 77 TIFAsome associated proteins that were enriched after *H. pylori* infection were identified by MS. The identified TIFA-binding proteins were used to generate PPI network graphs using STRING (Figure S3B). Interestingly, our data revealed that TIFAsomes contain several classical NF-κB key regulators, including TAB2 and TRAF2. Moreover, TRIM21, a factor involved in ubiquitination and proteasomal degradation, was enriched in this complex, consistent with its known role in NF-κB regulation (Oeckinghaus et al., 2011). Other prominent components of the TIFAsomes included the microtubule- and actin-associated proteins KIF11 and myosin IIA. Intriguingly, myosin IIA has been reported to be a target for ALPK1 in patients with gout (Lee et al., 2016). We also identified annexin A2, which was previously shown to interact with the p50 subunit of NF-κB to regulate p50 complex translocation into the nucleus (Jung et al., 2015). TRAF2 was chosen as an example for validating the MS data by immunoblotting and, evidently, co-precipitated with TIFA confirming the presence of TRAF2 in the *H. pylori*-induced TIFAsomes, consistent with the known interaction of TIFA and TRAF2 (Kanamori et al., 2002) (Figure S3C).

### CagA translocation and ALPK1-TIFA activation constitute independent pathways

In line with a number of previous studies (reviewed in Backert and Naumann (2010) p65 translocation, TAK1 activation and IL-8 mRNA upregulation in our system were dependent on the T4SS, but not on CagA (Figure S4A-S4C). To test whether activation of the ALPK1-TIFA axis itself also depends solely on T4SS function, we transfected AGS SIB02 cells with Tomato-TIFA for monitoring TIFAsome formation after infection (Figure 4A). As anticipated, TIFAsomes quickly formed after infection with wild type and CagA-deficient *H. pylori* but not the T4SS-deficient *cag*PAI mutant (Figure 4A). To test whether ALPK1 or TIFA in turn have an impact on CagA translocation, we determined CagA phosphorylation. Interestingly, CagA phosphorylation readily occurred in TIFA- or ALPK1-deficient cells (Figure 4B), suggesting that neither of the two factors is required for CagA translocation. To extend this analysis to other factors previously associated with CagA translocation, we generated a CRISPR/Cas9 knock out of the carcinoembryonic antigen-related cell adhesion molecule 1 (CEACAM1), recently identified as a receptor of HopQ (Javaheri et al., 2016; Koniger et al., 2016), an adhesin critical for T4SS function (Belogolova et al., 2013). In agreement with the proposed pleiotropic receptor recognition by HopQ, we observed partial dependency of both CagA translocation and NF-κB activation on the presence of CEACAM1 (Figure S4D-F). This places the branching point of the two processes, CagA translocation and activation of the ALPK1-TIFA axis, immediately downstream of the HopQ-CEACAM interaction.

**Fig. 4:**
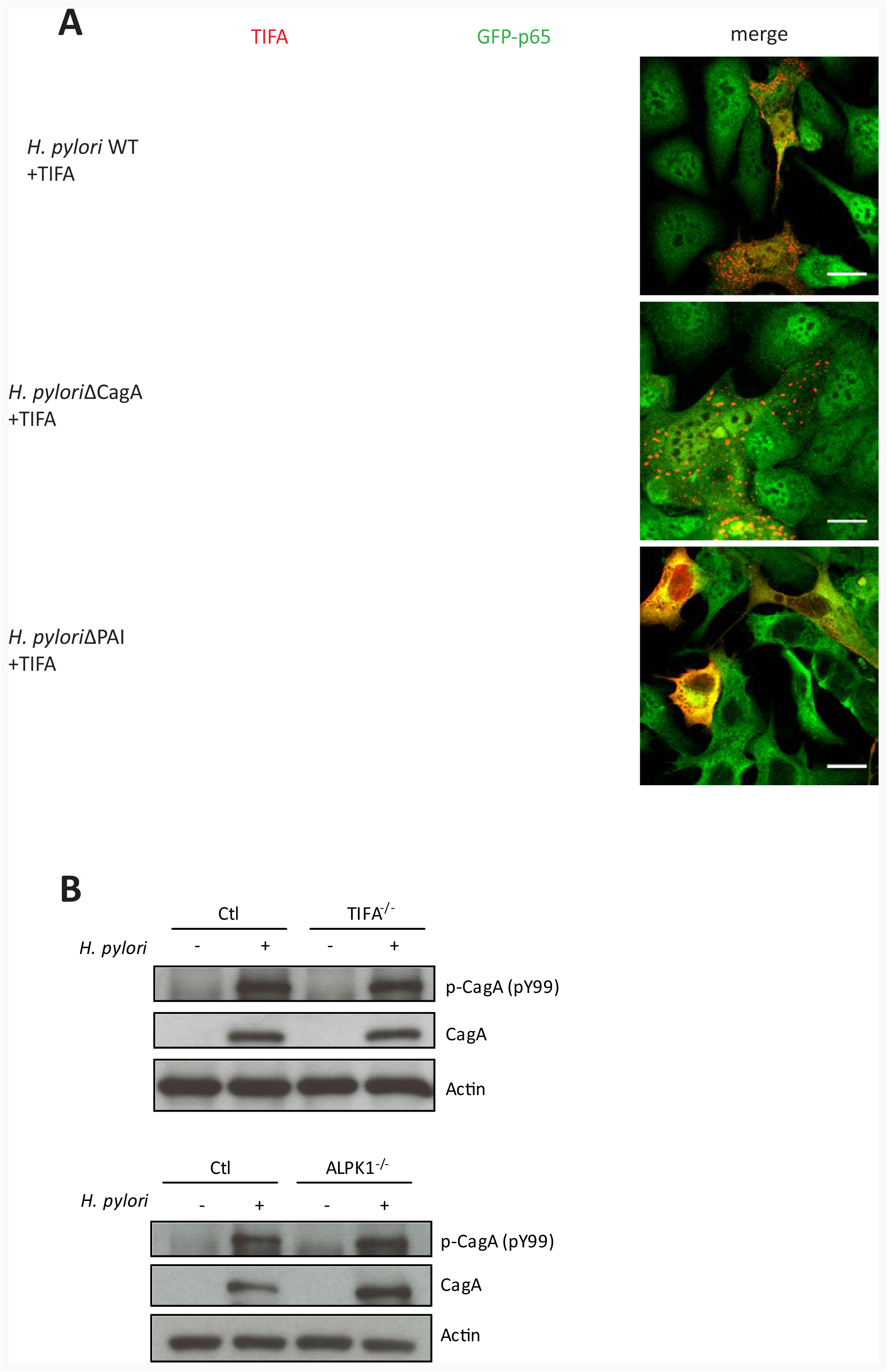
NF-κB activation and TIFAsome formation via ALPK1 and TIFA is independent of CagA translocation but dependent of a functional T4SS. (A) *H. pylori* WT and the *ΔcagA* but not the *ΔcagPAI* deletion mutant induce TIFAsome formation and NF-κB translocation. AGS SIB02 cells were transfected for 24 h with Tomato-TIFA (red) and then infected with *H. pylori* WT, *ΔcagA* or *ΔcagPAI* (MOI 100, 30 min). Scale bar: 20 μm. (B) CagA translocation and phosphorylation is not affected in cells depleted for TIFA or ALPK1 via CRISPR/Cas9. Cells were infected with *H. pylori* WT at MOI 100 and analyzed by Western blot at 3 h p.i. for tyrosine phosphorylation and total CagA. Blots are representative of two independent experiments.

### T4SS required for release of HBP and ALPK1-TIFA dependent NF-κB activation

We speculated that formation of TIFAsomes and subsequent activation of NF-κB could be triggered by a bacterial component secreted in a T4SS-dependent fashion. The pathogen-associated molecular pattern (PAMP) heptose-1,7-bisphosphate (HBP), which has recently been shown to activate TIFA (Gaudet et al., 2015; Milivojevic et al., 2017), appeared a likely candidate. HBP is a monosaccharide produced in the synthesis pathway of Gram-negative lipopolysaccharides (Valvano et al., 2002). We thus prepared bacterial lysates as previously described (Gaudet et al., 2015) from wild type *H. pylori*, as well as the corresponding Δ*cag*A, Δ*cag*L and Δ*cag*PAI mutant strains, and used them to transfect wild type, as well as TIFA- and ALPK1-deficient AGS SIB02 cells. qRT-PCR analysis showed that within 3 h IL-8 expression was upregulated in control cells transfected with lysates from both T4SS competent and T4SS deficient bacteria, as well as with lysate prepared from Gram-negative *N. gonorrhoeae* but not from Gram-positive *L. monocytogenes* (Figure 5A). By contrast, in ALPK1- or TIFA-deficient cells no increase in IL-8 expression was observed upon transfection of any lysates. Moreover, we observed TIFAsome formation and nuclear translocation of p65 in NF-κB reporter cells upon transfection of all *H. pylori*- (wild type, Δ*cag*A, Δ*cag*L and Δ*cag*PAI) and *N. gonorrhoeae*- but not *L. monocytogenes*-derived lysates (Figure 5B and data not shown). Translocation of p65 in NF-κB reporter cells transfected with *H. pylori* wild type lysate was also monitored by time-lapse video microscopy and, in contrast to the activation by live *H. pylori*, occurred in a non-synchronized fashion (Movie S2).

**Fig. 5.**
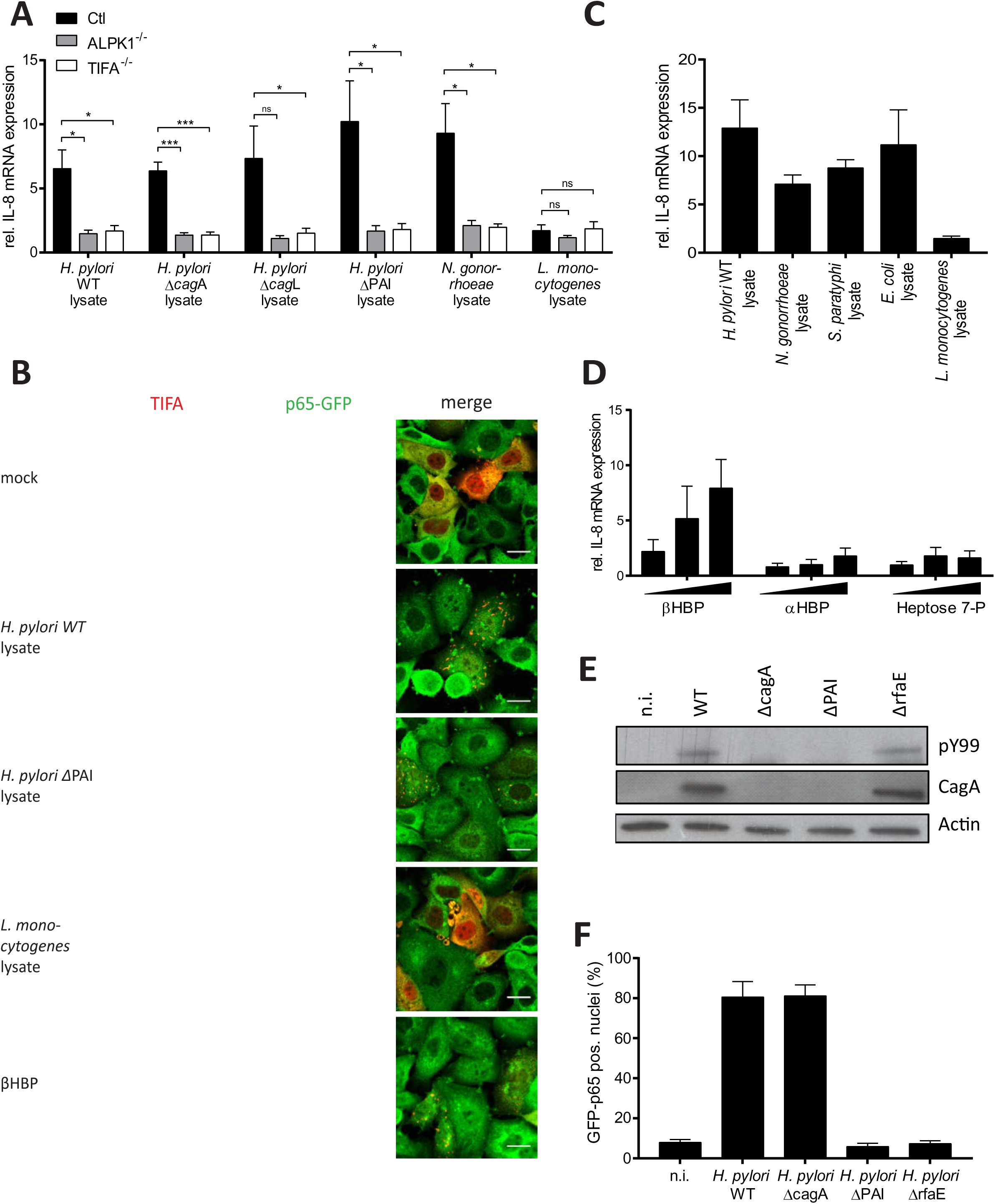
Bacterial lysates (HBP) trigger NF-κB activation dependent on ALPK1-TIFA axis. (A) Transfection with bacterial lysates derived from *H. pylori* T4SS competent (P12 WT and P12Δ*cag*A) and T4SS deficient strains induce IL-8 mRNA expression in AGS control cells but not in cells depleted of TIFA- or ALPK1 by CRISPR/Cas9. Bacterial lysates were prepared as described (Gaudet et al., 2015) and transfected into cells using lipofectamine. Levels of IL-8 mRNA were measured using qRT-PCR 3 h post transfection and normalized to mock transfected control cells. Data represent the mean ± SEM of three independent experiments. Statistical analysis was performed using Student’s t-test (*, *p* ≤ 0.05; **, *p* ≤ 0.001; ***, *p* ≤ 0.0001; ns = not significant). (B) TIFAsome formation induced by *H. pylori* lysates from wild type and T4SS mutant bacteria and by βHBP. AGS SIB02 cells were transfected for 24 h with Tomato-TIFA (red), then mock transfected or transfected with lysates derived from *H. pylori* WT, the ΔPAI mutant, *L. monocytogenes* or βHBP for 3 h, fixed and analyzed by microscopy. Scale bar: 20 μm. (C) Bacterial lysates from Gram-negative bacteria but not from Gram-positive *Listeria* induce IL-8 mRNA expression in AGS cells treated as in (A). Data represent the mean ± SEM of three independent experiments. (D) D-*Glycero*-β-D-*manno*-heptose 1,7-bisphosphate (βHBP) but not D-*glycero*-α-D-*manno*-heptose 1,7-bisphosphate (αHBP) or D-*glycero*-D-*manno*-heptose 7-phosphate (heptose 7-P) induce IL-8 mRNA expression in AGS cells 3 h post transfection, normalized to mock transfected control cells. (E) CagA phosphorylation can be detected in cells upon infection with a *H. pylori rfaE* mutant. Wild type cells and cells infected with *H. pylori* WT and Δ*rfaE* mutant at MOI 100 and analyzed by Western blot 3 h p.i. for tyrosine phosphorylation and total amounts of CagA. Blots are representative of two independent experiments. (F) NF-κB activation by *H. pylori* is dependent on expression of *rfaE*. AGS SIB02 cells were infected with wild type P12, Δ*rfaE*, Δ*cagA* and *ΔcagPAI H. pylori* (MOI 100, 30 min) and nuclear p65 translocation analyzed by automated microscopy. Unless otherwise indicated, data represent the mean ± SEM of two independent experiments. See also Movie S2 and S3.

IL-8 was also upregulated upon transfection with lysates from the Gram-negative *S. paratyphi* and *E. coli* (Figure 5C), further supporting the idea that the NF-κB-inducing factor is a molecule expressed specifically by Gram-negative bacteria such as HPB. Thus, we chemically synthesized D-*glycero*-β-D-*manno*-heptose 1,7-bisphosphate (βHBP), as well as its anomer D-*glycero*-α-D-*manno*-heptose 1,7-bisphosphate (αHBP), and the monophosphate D-*glycero*-D-*manno*-heptose 7-phosphate (heptose 7-P) (Figure S5A-C). Transfecting AGS cells with increasing amounts of βHBP, as compared to αHBP or heptose 7-P, induced IL-8 expression (Figure 5D) and formation of TIFAsomes (Figure 5B, lower panel), an observation further confirmed by time-lapse video microscopy of p65 translocation (Movie S3). This suggests NF-κB activation can be induced by translocation of the HBP β-anomer via the ALPK1-TIFA axis. Generation of a *H. pylori rfa*E knockout mutant corroborated the action of HBP as the specific inducer of NF-κB. RfaE catalyzes the phosphorylation of D-*glycero*-D-*manno*-heptose-7-phosphate at the C-1 position to generate HBP. Infection with this mutant still allowed for CagA translocation but not NF-κB activation (Fig. 5E and F). To demonstrate that the observed ALPK1-TIFA axis-dependent induction of NF-κB is not a cell line specific phenomenon, we tested gastric primary cells isolated from patient samples, expanded as organoids and seeded in 2D (Schlaermann et al., 2016) prior to transfection with recombinant TIFA. Consistent with our previous observations we found that *H. pylori* wild type, but not the *cag*PAI mutant, infection caused formation of TIFAsomes in these authentic gastric cells. This indicates that the ALPK1-TIFA signaling route is active in human primary cells (Figure 6).

**Fig. 6.**
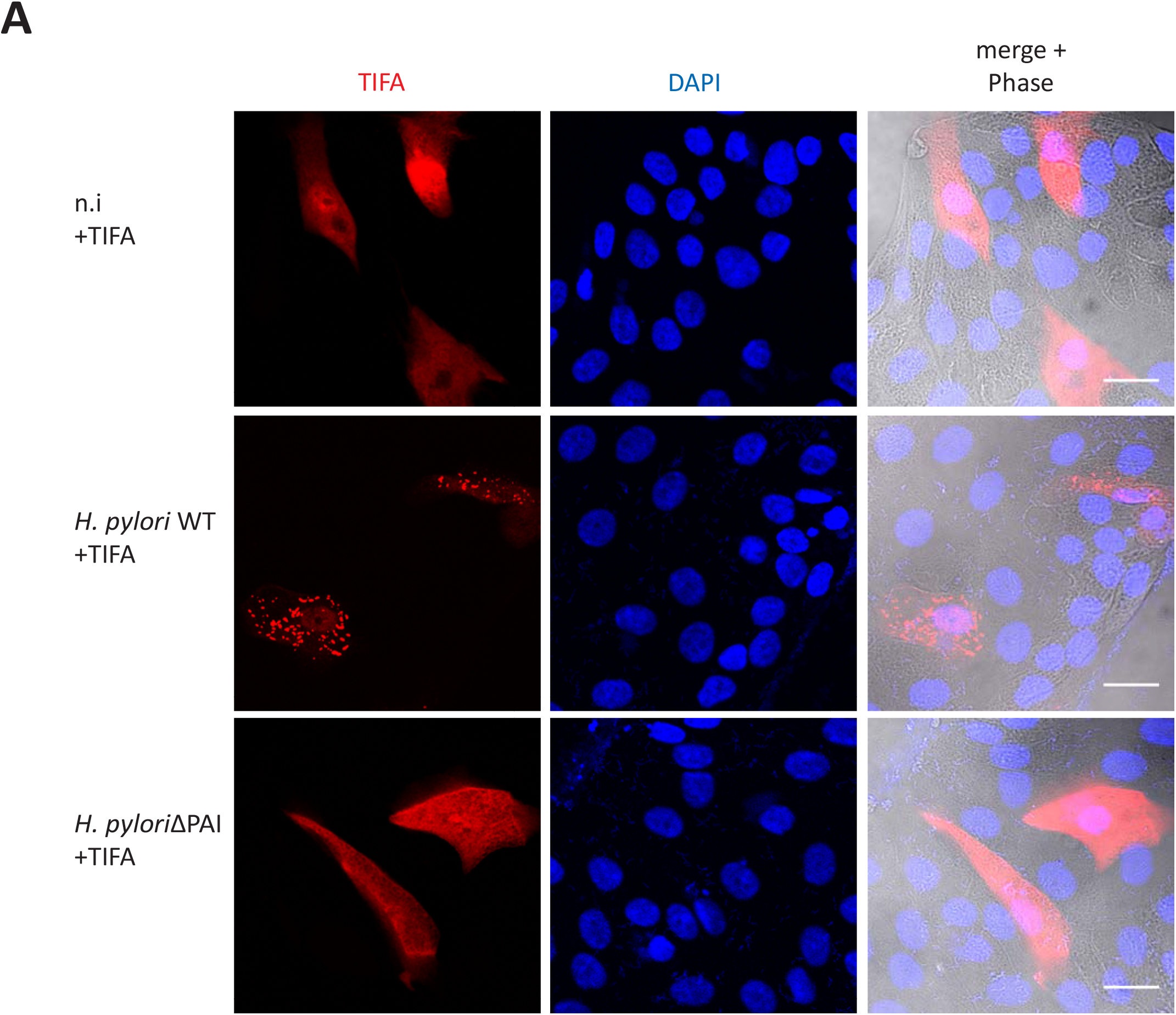
TIFAsome formation in primary gastric epithelial cells. Human gastric primary cells were grown as 2D monolayers as previously described (Schlaermann et al., 2016) and transfected with Tomato-TIFA for 24 h using FuGENE6. Cells were left uninfected or infected with *H. pylori* P12 WT and ΔPAI (MOI 100, 30 min) then fixed and counterstained for DNA (DAPI). As in AGS cells, infection with T4SS competent bacteria induced formation of TIFAsomes.

## Discussion

Discriminating pathogenic from non-pathogenic microbial traits is a prime function of the innate immune system and the epithelial barrier is equipped with exquisite sensory means to accomplish this task. We were interested in understanding the basis of the innate recognition of highly pathogenic strains of *H. pylori*, known to elicit particularly strong NF-κB-based immune reactions in the gastric mucosa depending on their *cag*PAI-encoded T4SS (Backert and Naumann, 2010; Koch et al., 2016). Our extensive screening program now provides a list of host factors involved in nuclear translocation of p65 selectively stimulated by *H. pylori* as compared to IL-1β and TNF-α. Of the main hits, ALPK1 identified in the initial kinome screen (Bartfeld, 2009) appeared to be furthest upstream in the pathway, triggered by the T4SS-dependent infusion of HBP to the host cell cytosol. ALPK1, either directly or indirectly, then causes TIFA phosphorylation and complex formation with an array of host factors, leading to activation of classical NF-κB signaling (Figure 7). This uncovers the HPB-ALPK1-TIFA axis as the key regulator of *H. pylori* T4SS-dependent NF-κB activation.

**Fig. 7.**
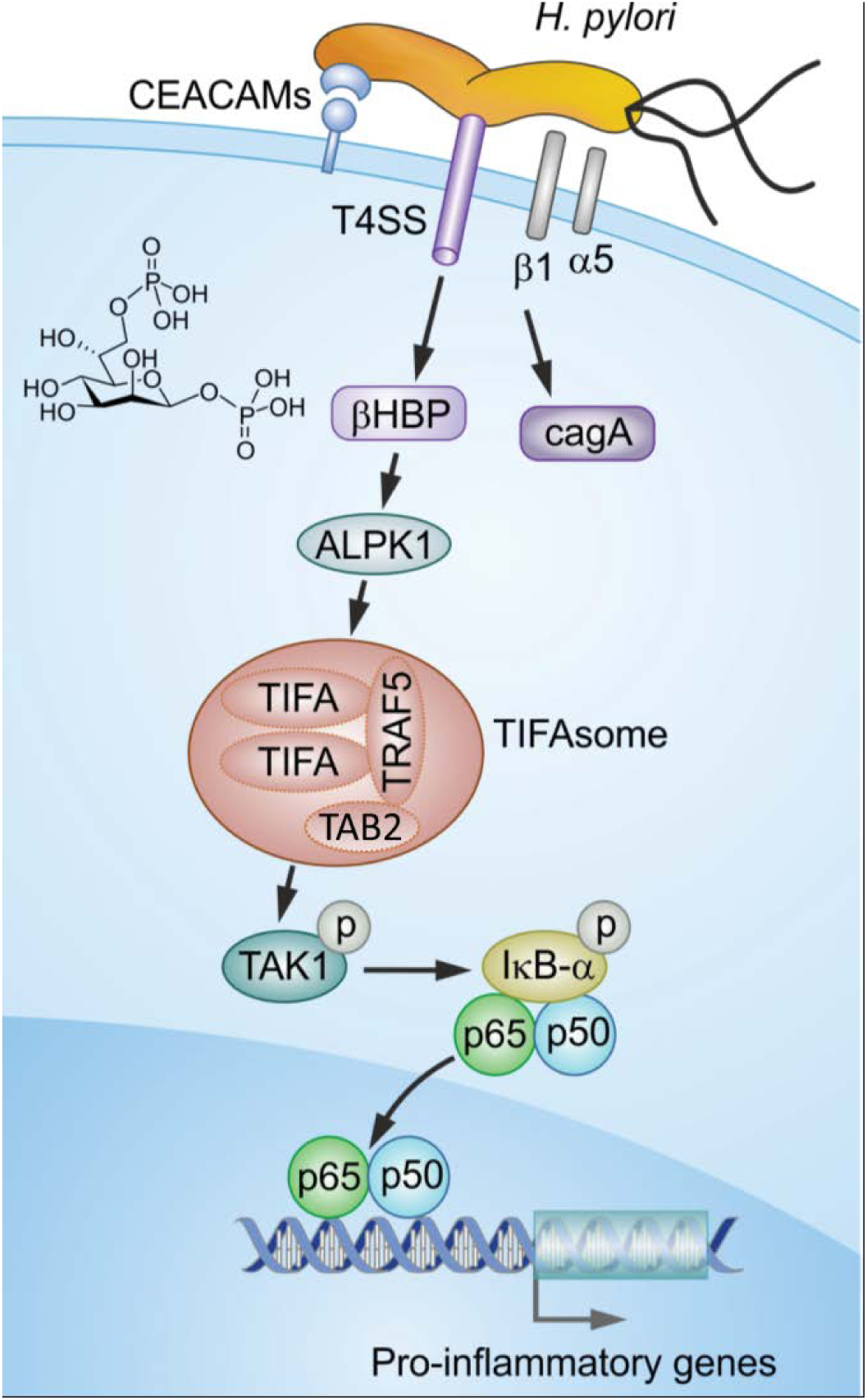
Schematic of *H. pylori* – host cell interaction leading to NF-κB pathway activation. *H. pylori* T4SS function depends on binding mediated by HopQ-CEACAMs and T4SS – α1β5 integrins and facilities the translocation of CagA and βHBP. βHBP in turn activates the ALPK1-TIFA-axis. cells 3 h post transfection, normalized to mock transfected control cells. (E) CagA phosphorylation can be detected in cells upon infection with a *H. pylori rfaE* mutant. Wild type cells and cells infected with *H. pylori* WT and Δ*rfaE* mutant at MOI 100 and analyzed by Western blot 3 h p.i. for tyrosine phosphorylation and total amounts of CagA. Blots are representative of two independent experiments. (F) NF-κB activation by *H. pylori* is dependent on expression of *rfaE*. AGS SIB02 cells were infected with wild type P12, Δ*rfaE*, Δ*cagA* and Δ*cagPAI H. pylori* (MOI 100, 30 min) and nuclear p65 translocation analyzed by automated microscopy. Unless otherwise indicated, data represent the mean ± SEM of two independent experiments. See also Movie S2 and S3. See also Figure S5.

ALPK1 is a member of the atypical kinase family alpha kinases that recognize phosphorylation sites within α-helices (Ryazanov et al., 1999). Until recently, the only substrates suggested for ALPK1 were myosin IA and IIA (Heine et al., 2005; Lee et al., 2016). Indeed, we found myosin IIA to be part of the TIFAsomes but whether it is phosphorylated by ALPK1 or plays a role in the activation of innate immunity during *H. pylori* infection has to be elucidated. A recent report also identified ALPK1 as necessary for TIFA oligomerization after infection with *S. flexneri* (Milivojevic et al., 2017), connecting it to the innate immune response. A role for ALPK1 and TIFA in inflammation was first predicted from an analysis of major genetic susceptibility loci in the experimental IL10^-/-^ mouse model of inflammatory bowel disease (Bleich et al., 2010). ALPK1 was later found to be upregulated in patients with gout and to modulate the expression of inflammatory cytokines after the addition of ureate crystals in an experimental gout model (Wang et al., 2011). The role of ALPK1 in these inflammatory scenarios is still unknown, but the importance in innate immune sensing of *S. flexneri* (Milivojevic et al., 2017) together with the data presented here point to a general role of ALPK1 in HBP sensing, making it a key regulator of inflammation.

TIFA is the direct neighbor of ALPK1 on human chromosome four. It is a ubiquitously expressed cytoplasmic protein first identified as a TRAF2 binding protein that activates the NF-κB pathway (Kanamori et al., 2002). TIFA phosphorylation at T9 promotes NF-κB activation through selfoligomerization and interaction with TRAF (Huang et al., 2012) and has recently been shown to be triggered by HBP (Gaudet et al., 2015) or by oxidative stress via Akt kinase (Lin et al., 2016). The kinase activity of ALPK1 is required for TIFA oligomerization after infection *with S. flexneri* (Milivojevic et al., 2017) and *H. pylori*, as we show here. However, whether ALPK1 directly phosphorylates TIFA at T9, which is not within the α-helix classically recognized by atypical kinases, remains to be resolved.

Both our RNAi screening and mass spectrometry analyses point towards a complex array of host factors involved in NF-κB activation by *H. pylori* whose functions remain largely unknown. However, amongst the positive regulators in the RNAi screens we identified HCF1, ASL2, and MLL4 as important for ALPK1 transcriptional activation. Future downstream target analysis might provide insights into genetic programs co-regulated with ALPK1. Specificity for *H. pylori* was also seen for two acetyltransferases, ARD1 and NARG1. Moreover, immune precipitation of TIFA revealed a large protein complex consisting of factors that likely integrate signals upstream from ALPK1 and guide them towards the more classical NF-κB route. In fact, TIFAsomes contained a number of core NF-κB pathway components, including TRAF2 and TAB2, as well as many more, whose contribution to NF-κB signaling (and potentially also additional pathways) awaits future analysis. Interestingly, ALPK1 itself did not co-localize with TIFAsomes, as evidenced by both immune precipitation/mass spectrometry and immunofluorescence of transfected cells, pointing towards a transient or indirect phosphorylation mechanism of this kinase.

As the *H. pylori* component responsible for NF-κB activation, we here identify the small molecule metabolite βHBP, which is a precursor of LPS previously recognized as a PAMP of several Gram-negative bacteria, including *N. gonorrhoeae* (Gaudet et al., 2015) and *S. flexneri* (Milivojevic et al., 2017). We demonstrate the strict dependency of NF-κB activation on the pathogen’s functional T4SS, suggesting that it delivers βHBP to the inside of host cells. Accordingly, mutants of not only the T4SS, but also the respective LPS biosynthetic pathway (*rfa*E), fail to induce NF-κB and downstream IL-8. Consistently, we found no evidence in support of the previously proposed function of CagA in NF-κB activation (Lamb et al., 2009), nor of the proposed role of small peptidoglycans supposedly translocated by *H. pylori* (Viala et al., 2004). Moreover, no significant hits in our screens pointed to a role of NOD1, NOD2, or respective downstream factors (Lipinski et al., 2012; Warner et al., 2013; Yeretssian et al., 2011).

The T4SS machinery is thought to be activated by the interaction of certain cagPAI-encoded ligands (CagL, CagA, CagY, CagI) with members of the integrin family (Kwok et al., 2007, Jiménez-Soto et al., 2009, Rohde et al., 2003). In addition, we have identified HopQ as a further factor critical in this process (Belogolova et al., 2013) and subsequent work in other laboratories has shown that HopQ interacts with members of the CEACAM family of receptors (Koniger et al., 2016, Javaheri et al., 2016). Both types of interaction appear to be crucial for translocation of the CagA effector protein (Belogolova et al., 2013, Oleastro and Ménard, 2013). Interestingly, activation of the ALPK1-TIFA-NF-κB axis is independent of CagA, and the *rfa*E mutant still supports CagA translocation. These findings mechanistically separate CagA translocation from βHBP delivery, although both depend on the functional integrity of T4SS. While we show that this striking pathway functions readily in authentic normal human gastric primary cells, and that TIFA activation occurs in the very first minutes of *H. pylori* infection of the epithelium, it constitutes the initial trigger of a broader inflammatory cascade initiated by this pathogen, to which additional autocrine and paracrine cellular pathways contribute (Backert et al., 2016; Velin et al., 2016). Here, we have identified a novel mechanistic principle of how pathogenic *H. pylori* bacteria are recognized by innate immune defense at the stage of initial epithelial cell contact. This finding will guide future work towards a better understanding of pathogen defense and gastric pathogenesis.

## Experimental Procedures

### Ethical permissions

For Primary cell preparation, gastric tissue samples from individuals undergoing gastrectomy or sleeve resection were obtained from the Clinics for General, Visceral and Transplant Surgery, and the Center of Bariatric and Metabolic Surgery, Charité University Medicine, Berlin, Germany. Usage of the pseudonymized samples for experimental purposes was approved by the ethics committee of the Charité University Medicine, Berlin (EA1/058/11 and EA1/129/12).

### Bacterial strains and cultivation

The following *H. pylori* strains were used in this study: P1 and P12 wild type (strain collection no. P213 and P511) and the mutant strains P12Δ*cag*PAI, P12Δ*cag*A and P12Δ*cag*L and P12*ΔrfaE* (strain collection no. P387, P378, P454 and 588). *H. pylori* cultivation was carried out as described before (Backert et al., 2000). If not otherwise mentioned MOI 100 was used.

### Cell Culture

AGS cells stably transfected with a p65-GFP construct (AGS SIB02, NF-κB reporter) (Bartfeld et al., 2010), AGS SIB02 CRISPR-Cas9 control and knockout cells (AGS STZ001, AGS STZ003 and AGS STZ004) and the parental AGS cells (ATCC CRL 1739, human gastric adenocarcinoma epithelial cell line) were cultivated in RPMI medium (Gibco) supplemented with 10% heat inactivated fetal calf serum and 2 mM L-glutamate. Cultures were kept at 37°C in a humidified atmosphere with 5% CO_2_. AGS cell infections were carried out after serum starvation in serum free medium.

### Primary cell culture

Culture of gastric primary glands was performed as described by Schlaermann et al. (2016) Briefly, cells from freshly isolated glands were seeded in Matrigel for the formation of organoids and expanded. Cells were sheared every 10 days and for experimental use seeded on collagen coated coverslips and grown for 1-2 days in 2D medium.

### p65 translocation assay

AGS SIB02 cells were seeded into 12-well plates and infected with *H. pylori* (MOI 100, 30min) or stimulated by IL-1β (10 ng/ml, Peprotech) or TNF-α (20 ng/ml, Peprotech) for 30 min under serum-free conditions, fixed with ice-cold Methanol, nuclei counterstained with Hoechst 33342 (2 μg/ml, Molecular Probes). Images were acquired using automated microscopy (Olympus) NF-κB translocation was analyzed using an automated cell-based assay.

### Screening Procedure

For the kinome screen we used the Qiagen “Human Kinase siRNA Set”, containing 1,292 siRNAs targeting 646 kinase and kinase-associated genes. To reduce sample size, two siRNAs for each gene were pooled and tested in six conditions: *H. pylori* (45 min and 90 min), TNFα (30 min and 75 min) and IL 1β (45 min and 90 min) using four independent biological replicates. AGS SIB02 cells were seeded onto 96-well plates and transfected using the RNAiFect Transfection Kit (Qiagen). One day after transfection, cells were split into new wells (96-well-plate or 12-well-plate according to experimental setting). Experiments were conducted after a minimum of 60 h to allow reduction of target protein levels. Data was normalized (either using plate-median or, in comparison, z-score) and significance was assessed with Welch’s t-test. Primary hits were identified combining statistical significance (p-value ≤0.05). For the validation screen the 5% top candidates were taken into account for further analysis and an additional 4 siRNAs were tested and hits were confirmed with at least two siRNAs each with a z-score of ≤-1 or ≥1.

For the primary screen the Qiagen Human Druggable Genome siRNA Set V2.0 Library containing 4 siRNAs per gene and the Qiagen Human Genome 1.0 Library containing 2 siRNAs per gene were used. Altogether 24,000 genes were targeted by 62,000 individual siRNAs. For the validation screen 4 individual siRNAs were purchased from Qiagen. AGS SIB02 cells were seeded into 384-well plates, siRNAs were transfected with HiperFect transfection reagent (Qiagen), 72 h post transfection cells were stimulated for 45 min by infecting with *H. pylori* (MOI 100) or in the validation screen cells were additionally induced by IL-1β (10 ng/ml, Miltenyi Biotec.) or TNF-α (10 ng/ml, Becton Dickinson) for 45 min under serum-free conditions, fixed with ice-cold methanol, nuclei counterstained with Hoechst 33342 (2 μg/ml, Sigma Aldrich). Images were acquired using automated microscopy (Olympus) NF-κB translocation was analysed using an automated cell-based assay as recently described (Bartfeld et al., 2010). In short, cell nuclei were detected and the surrounding cytoplasmic area set using image analysis software (Scan^R, Olympus) following quantification of translocation of p65-GFP. Experiments were performed in triplicates. Data analysis primary screen: First, plate normalization was performed using CellHTS2, then Redundant siRNA Analysis (RSA) was used for defining the hit candidates. Positive regulators are defined by an RSA-score ≥−2, negative regulators by an RSA-score of ≥+2. Hit validation screen was performed for 300 selected primary hit candidates (200 positive and 100 negative regulators) using 4 siRNAs per gene. Cells were transfected for 72 h and NF-κB activated with either *H. pylori* infection, TNF-α or IL-1β. Cells were fixed after 45 min and p65-GFP translocation automatically quantified. Data were analyzed using CellHTS2. Hit genes were defined according to their Z-score: <−1 for positive and >1 for negative regulators. As a positive control for all screening experiments the combinatorial knockdown of IKKα and β was used.

### Immunoblotting

Cells were directly lysed in 2x Laemmli buffer, separated by 10% SDS-PAGE, transferred to PDVF membranes, blocked in TBS buffer supplemented with 0.1% Tween 20 and 5% milk, and probed against primary antibodies at 4°C overnight. Membranes were probed with matching secondary horseradish peroxidase-conjugated antibodies (Amersham, 1:3000) and detected with ECL reagent (Perkin Elmer). Primary antibodies: Anti IκBα (44D4) #4812, p-TAK1 (Thr 184/187) #4508 and p-IκBα (Ser32) (14D4) #2859, pNF-κB p65 (Ser536) #3033 from Cell Signaling; anti phospho-tyrosine 99 (pY99) sc-7020, anti CagA (b300) sc-25766, and β-actin A5441 from Sigma Aldrich.

### qRT-PCR

qRT-PCR was performed using Power SYBR Green RNA-to-C_T_ 1-Step Kit (Applied Biosystems) according to the manufacturer’s recommendations using the following primers: GAPDH 5′-GGTATCGTGGAAGGACTCATGAC-3′ and 5′-ATGCCAGTGAGCTTCCCGTTCAG-3′, IL-8 5′-ACACTGCGCCAACACAGAAAT-3′ and 5′-ATTGCATCTGGCAACCCTACA-3′, 5′-TGGTAAACCGTCATCTGGAG-3′ and 5′-GAGTTCACTGACTCCCCAGC -3′. ALPK1 5′-CACCAAGAACACAATAGCCG-3′ 5′-ACCTGAAGGATGTGATTGGC-3′

### Plasmids

For overexpression of *N*-terminal wild type Myc-Flag-TIFA we used full length human TIFA cDNA from Origene #RC204357 were we introduced a silent TGG to TCG point mutation encoding L103 to destroy the PAM site preventing CAS9 cutting of TIFA when overexpressed in TIFA deficient cells (strain collection no. pMW930). Mutation of the PAM site and TIFA T9A mutation (strain collection no. pMW931) were introduced by using PfuTurbo DNA Polymerase (Agilent Technologies) complying with the protocol of Stratagene’s single-site QuikChange *Site-Directed Mutagenesis* Kit. For generating *N*-terminal Tomato-TIFA, TIFA from pMW930 was amplified and cloned into ptdTomato N1 (modified from Clonetech #6085-1) (strain collection no. pSTZ006). Human ALPK1 was amplified and cloned into pLenti-*N*-Myc-DDK (strain collection no. pMW909).

Myc-Flag-ALPK1 5′- CTGCCGCCGCGATCGCCATGAATAATCAAAAAGTGGTAGC-3′ and 5′- TTGCGGCCGCGTGCATGGTTTCTCCATTGAAG-3′, Tomato-TIFA 5′- CGGCTAGCCGCCATGACCAGTTTTGAAG -3′ and 5′- CGGAATTCCGGTTGACTCATTTTCATCCATTTCT -3′

### Confocal microscopy

AGS, AGS SIB02 or primary cells were seeded on poly-l-lysine or collagen coated glass coverslips and transfected for 24h with recombinant ALPK1, recombinant TIFA or its functional mutants using FuGENE6 (Promega). Cells were left uninfected or infected with *H. pylori* (MOI 100), treated with TNF-α (20 ng/ml) or transfected for 3 h with bacterial lysates using lipofectamine 2000 (Thermo Fisher Scientific) under serum free conditions. For visualization of MYC-tagged protein, cells were fixed with 4 % PFA, blocked with 3% BSA, 5%FCS, 0.3Tritx100 for 1 h at RT and incubated with primary antibodies (anti-MYC (CST, Cat. No 9B11, 1:6000), anti-tubulin (Abcam, Cat. No. ab 6160, 1:100)) overnight at 4°C. Secondary staining was performed using fluorescently coupled antibodies (donkey anti rabbit-Cy3 (1:100, Cat. No. 711-166-020), goat anti rat-Dylight^®^ 649 (1:100 Cat. No. 112-486-062), DAPI (1:300, Roche, Cat. No. H1840-10)). Samples were mounted with Moviol, analyzed by laser scanning microscopy using a Leica SP8. Images were processed using ImageJ.

### Bacterial lysate preparation and transfection

Bacterial lysates were prepared as described in Gaudet et al., 2015 (Gaudet et al., 2015). In short, *H. pylori* and *N. gonorrhoeae* were grown overnight on GC agar plates, *E. coli* was grown overnight on LB agar plates, *L. monocytogenes* was grown overnight in brain heart infusion medium and *S. paratyphi* was grown overnight in LB media. Bacteria from overnight cultures were collected in PBS and resuspended in water at OD_600_ 1 and boiled for 15 min at 400 rpm shaking. Bacterial debris was removed by centrifuged at 4000 × g for 3 min and the supernatant treated with 10 μg/mL RNAse A (New England Biolabs), 1 U/mL DNAse 1 (Qiagen) and 100 μg/mL ProteinaseK (New England Biolabs) for 5 min at RT. Samples were boiled again for 5 min and passed through a 0,22 μm filter. For lysate transfection of cells in a 24-well format, 5 μl lysate was mixed with 5 μl lipofectamine 2000 (Thermo Fisher Scientific) and 30 μl OptiMEM (Gibco), incubated for 30 min at RT and added to AGS cells at 70% confluence.

### Statistical analysis

All graphs were prepared using GraphPad Prism 7 software (GraphPad Software Inc., CA, USA). Data are presented as mean ± SEM. Data was considered significant if p<0.05.

## Additional Supplemental Files

### Table S1 Results of kinome screen

Names of genes are listed alphabetically with gene identification number (Gene ID). accession numbers (Accession #), mean numbers of cells counted in four pictures of a well (Cell #) and z-scores, which give indication of the strength of the effect. Negative z-scores indicate inhibition and positive z-scores indicate promotion of p65-translocation. In the primary screen as well as in the hit validation, each siRNA was tested in six conditions: H1 = *H. pylori* 45 min; H2 = *H pylori* 90 min; T1 = TNFα 30 min; T2 = TNFα 75 min; I1 = IL-1β 45 min; I2 = IL-1β 90 min. In the primary screen. two siRNAs per gene were pooled and the mean z-score of four experiments using this pool is shown. To identify primary hits. bold colored font marks the z-score in the condition in which the gene scored as a primary hit during the primary screen (red for inhibition, blue for promotion of p65-translocation). In the hit validation four siRNAs per gene were tested separately and for each siRNA. a mean z-score of four experiments was calculated. Here, the sum of the z-scores of four siRNAs is shown. To identify z-score sums composed of at least two effective siRNAs with z-scores of ≤−1 or ≥1. Bold colored font marks the z-score-sum in the condition in which at least two siRNAs were effective (red/blue as above). Colored cells and gene names in bold colored font indicate confirmed hits, defined as genes where at least two siRNAs were effective in the hit validation in the same condition as it was selected for in the screen (red/blue as above).

### Table S2 Results of primary and validation screen

Primary screen: 24,000 genes were silenced using Qiagen siRNA libraries. NF-κB activation was induced with *H. pylori*. Experiments were done in triplicates. CellHTS2 and Redundant siRNA Analysis (RSA) was used for defining the hit candidates. According to the corresponding RSA-score, 235 candidates were found to positively regulate and 112 factors to negatively regulate *H. pylori* NF-κB pathway. Validation screen: 300 candidate genes (200 positive and 100 negative regulators from the primary screen) were tested. *H. pylori* and in addition IL-1β and TNFα were used as inducers for NF-κB activation. Data analysis based on CellHTS2. The Venn diagram shows the validated factors. Apart from general factors functioning for all inducers, 21 positive regulators and 24 negative regulators unique for the *H. pylori* NF-κB pathway were found.

### Table S3 Results of mass spectrometry analysis of protein–protein interactions of TIFAsomes

Composition of 78 candidate proteins identified upon H. pylori infection as analyzed by UHPLC/MS/MS.

### Movie S1 *H. pylori* infection activates NF-κB via TIFAsome formation

AGS SIB02 TIFA knockout cells were seeded on 8μ-well dishes (Ibidi) and transfected with pdtTomato-TIFA. Cells were infected with *H. pylori* (MOI 100) and imaged on a Zeiss Axiovert 200M wide field microscope with an incubator at 37°C and 5% CO_2_ using a Hamamatsu Orca camera. Images for Tomato-TIFA and DIC were recorded every 15 s. The setup was controlled by the Volocity software (Perkin Elmer) which was also used to generate the video.

### Movie S2 Translocation of p65 in NF-κB reporter cells transfected with *H. pylori* lysate

AGS SIB02 cells were seeded on 8μ-well dishes (Ibidi) and transfected with lysates from *H. pylori*. Live cell imaging was performed using a Leica TCS-SP5 inverted confocal microscope with an incubator at 37°C and 5% CO_2_. Frames for GFP and DIC were recorded every 30 s and merged into a video using Volocity software (Perkin Elmer).

### Movie S3 βHBP induces p65 translocation

AGS SIB02 cells were seeded on 8μ-well dishes (Ibidi). Each well of the plate was transfected with a defined concentration of HPB. Wells were imaged in succession by constantly moving the stage from well to well. Images were recorded on a Zeiss Axiovert 200M wide field microscope with an incubator at 37°C and 5% CO_2_ using a Hamamatsu Orca camera. Images for eGFP and DIC were recorded every 2 mins. The setup was controlled by the Volocity software (Perkin Elmer) which was also used to generate the videos.

## Author contributions

S.B., N.M. performed kinome screen and revealed ALPK1; J.L., N.M. carried out the genome-wide screen; A.M., K.P.P., M.R. performed bioinformatics analyses; M.K., F.G. M.R. recognized factors including TIFA; C.R., M.R., S.B., O.S., M.N. performed NF-kB pathway analysis; S.Z. validated factors using CRISPR/Cas9; S.Z., M.A.Z., L.P. characterized HBP-ALPK1-TIFA interactions and performed cell assays; M.S. generated genetic constructs; A.Z., P.K. synthesized HBP; M.S. performed mass spectrometry; V.B., M.A.Z., L.P. confocal microscopy; T.F.M. conceived and supervised the project; S.Z., M.A.Z., L.P., S.B., T.F.M. wrote manuscript;

## Acknowledgements

We thank Jörg Angermann, Elke Ziska, Jan-David Manntz, Kathrin Lättig and Isabella Gravenstein for technical assistance, Rike Zietlow for editorial assistance. We are very thankful to Ralf Jacob and Katharina Cramm-Behrens for ALPK1 expression construct.

